# A New Serological Autoantibody Signature Associated with Multiple Sclerosis

**DOI:** 10.1101/2025.04.23.650160

**Authors:** Houari B. Abdesselem, Alberto D. Van Benthem, Ilham Bensmail, Israa E Elbashir, Ti-Myen Tan, Bermet Abylova, Lay Koon, Diana Anuar, Jonathan M. Blackburn, Rayaz A. Malik, Ioannis N. Petropoulos

**Affiliations:** Proteomics Core Facility, Hamad Bin Khalifa University (HBKU), Qatar Foundation, Doha, Qatar; College of Health and Life Sciences (CHLS), Hamad Bin Khalifa University (HBKU), Qatar Foundation, Doha, Qatar; Qatar Biomedical Research Institute, Hamad Bin Khalifa University (HBKU), Qatar Foundation, Doha, Qatar; Department of Integrative Biomedical Sciences, Faculty of Health Sciences, University of Cape Town, South Africa; Sengenics Corporation, Level M, Plaza Zurich, Damansara Heights, Kuala Lumpur 50490, Malaysia; Institute of Infectious Disease and Molecular Medicine, Faculty of Health Sciences, University of Cape Town, South Africa; Weill Cornell Medicine - Qatar, Doha, Qatar

**Keywords:** Multiple Sclerosis, Immunoproteomics, autoantibodies, SOX-10, MX1, ISG20, viral infection

## Abstract

The role of autoantibodies in the pathogenesis of Multiple Sclerosis (MS) remains incompletely understood. In this study, we analyzed serum samples from a cohort of MS patients in Qatar using high-throughput KoRectly Expressed (KREX) immunome protein-array technology. Compared to healthy controls, MS patients showed significantly altered autoantibody responses to 129 proteins, with a notable enrichment in autoantibodies targeting antiviral immune response-related proteins. Machine learning analysis identified a distinct molecular signature comprising 17 differentially expressed autoantibodies, including those against MX1, ISG20, MAX, SUFU, NR1H2, HMGN5, and EPHA10. Among these, autoantibodies against MX1-a key effector in the interferon-alpha/beta signaling pathway-showed the most pronounced increase, with nearly a threefold elevation in MS patients. While MX1 has previously been implicated in MS, this is the first report of autoantibody reactivity against the protein, suggesting a potential role in disease onset and progression. These findings support a link between antiviral immune responses and MS pathophysiology and offer a promising blood-based autoantibody signature that could inform future diagnostic and therapeutic strategies.

## Introduction

According to the Global Atlas of Multiple Sclerosis (MS), approximately 2.9 million individuals worldwide are currently living with the disease, with its prevalence continuing to rise^1^. Although the precise cause of MS remains unknown, a growing body of evidence supports an autoimmune origin^2, 3, 4^. The etiology of MS is multifactorial, involving genetic predisposition, environmental factors—particularly viral infections—and the subsequent development of an aberrant immune response directed against components of the central nervous system (CNS). MS is characterized as a chronic, immune-mediated, inflammatory, and demyelinating disorder of the CNS and is among the leading causes of neurological disability in young adults. Its pathological hallmarks include demyelination, axonal loss, and neurodegeneration^5^. Approximately 85% of patients are initially diagnosed with relapsing–remitting MS (RRMS)^6, 7^, which involves acute episodes of neurological dysfunction followed by partial or complete recovery. Over time, many RRMS cases transition into secondary progressive MS (SPMS), marked by a gradual and irreversible accumulation of disability^8^.

Since the FDA’s approval of interferon beta-1b in 1993—the first disease-modifying therapy (DMT) for MS—there has been significant progress in the development of immunomodulatory treatments, reducing inflammation and nerve damage ^9, 10^. By 2020, nine distinct classes of DMTs had been approved, offering expanded options for RRMS management^11^. Nevertheless, these therapies do not cure the disease, nor can they reverse existing neurological damage. Many patients continue to experience disease progression and long-term disability despite treatment^12, 8^.

A major challenge in MS research and clinical management is the lack of reliable biomarkers to predict disease onset, monitor progression, or guide therapeutic decisions. Despite extensive efforts, biomarker discovery has yielded limited and often inconclusive results. The development of robust biomarkers would enhance our understanding of MS pathophysiology, support patient stratification, improve therapeutic outcomes, and ultimately aid in the development of more effective, personalized interventions.

In this study, we address this unmet need using a high-throughput biomarker discovery platform to investigate the humoral immune response in MS. By profiling autoantibody responses to a comprehensive panel of 1,600 human proteins—including kinases, cytokines, growth factors, interleukins, transcription factors, and signaling molecules—we aim to uncover novel autoantibodies associated with disease activity, severity, and progression. These findings may provide critical insights into MS pathogenesis and support the development of blood-based biomarkers with diagnostic and therapeutic potential.

## Methods

### Study design, samples collection and processing

We used blood samples and clinical data of patients from MS cohort to conduct a comprehenisve anlaysis of autoantibodies using novel KREX technology. The MS cohort included Thirty-nine (n=39) patients from Qatar who were enrolled at Hamad Medical Corporation hospitals. All recruited patients had confirmed MS pathology. Peripheral blood was collected and processed into serum, which were stored at –80°C, until further analysis (**Figure 1**). Age and gender matched healthy volunteers (n=38) with no medical history. All participants (patients and controls) provided written informed consent prior to enrolment in the study.

**Figure 1.**
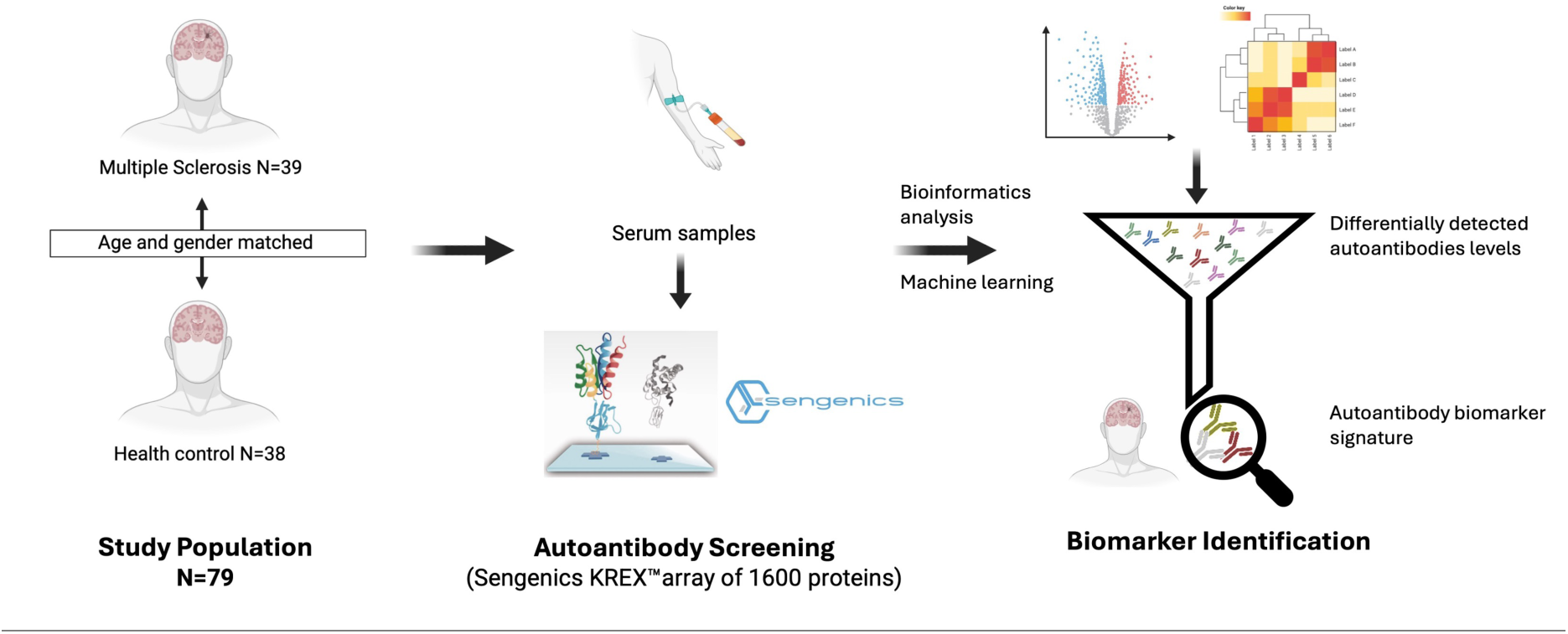

### Sengenics assay description and data pre-processing

MS cohort samples were processed at HBKU Proteomics Core Lab for KoRectly Expressed (KREX) immunoproteomics. Serum samples were analyzed for antigen-specific autoantibodies using Immunome protein arrays from Sengenics company, developed using KREX technology to provide a high-throughput immunoassay based on correctly folded, full length and functional recombinant human proteins expressed in insect cells, thereby displaying a full repertoire continuous and discontinuous epitopes for autoantibody binding^13, 14^. The Immunome arrays contain more than 1,600 human antigens, enriched for kinases, transcription factors, signaling molecules, cytokines, interleukins, chemokines, as well as known autoimmune antigens^15^.

Samples were diluted in Serum Albumin Buffer (SAB) at optimized dilution (400-fold dilution). Microarray slides were prepared in four-well plates slide. Samples including controls were randomized and applied to the microarray slides for 1 hour and samples’ IgGs were then detected by secondary Cy3-labeled IgG antibodies. Slides were scanned at a fixed gain setting using the Agilent G4600AD fluorescence microarray scanner generating a 16-bit TIFF file. A visual quality control check was conducted and any array showing spot merging or other artefacts were re-assayed. A GAL (GenePix Array List) file containing information regarding the location and identity of all probed spots was used to aid with image analysis. Automatic extraction and quantification of each spot was performed using GenePix Pro 7 software (Molecular Devices) yielding the median foreground and local background pixel intensities for each spot.

Biotinylated human IgG (detected by fluorescently labelled secondary antibody) and biotinylated human anti-IgG (detected only when serum is added to the slide) were used as positive controls to assess assay integrity. Extrapolated data was then filtered, normalized and transformed as follows. The median background pixel intensity (RFU) was subtracted from the median foreground pixel intensity (RFU) for each antigen to give the median net intensity per spot (RFU). CVs were calculated for each antigen based on the quadruplicate technical replica spots for each antigen on a given array. Antigens with CVs above 20% were flagged and outlier spots were removed, provided that at least two valid values remain. Net intensity value for each antigen in a given sample was calculated as the mean of the net intensity values for technical replica spots on that array. The data was normalized across replica arrays based on the Cy3-BSA controls as previously described^16^.

### Bioinformatics data analysis

Bioinformatics pipeline was implemented using R programming packages (**R version** 4.4.1). Using Sengenics data, differential expression analysis has been performed between MS and control groups using the LIMMA package (https://www.bioconductor.org/packages/release/bioc/html/ limma.html). Thresholds of p-value < 0.05 and FC > 1.25 were used to identify differentially expressed proteins. Raw p-values (not adjusted) were considered. Visualization of significant changes in protein expression levels was achieved through the creation of Heatmap (pheatmap package), and Box plots (gglot2), and Volcano plots using the EnhancedVolcano package (https://bioconductor.org/packages/release/bioc/html/Enhanced Volcano.html). Machine learning (ML) modeling has been conducted to identify minimal set of biomarkers that are specific to MS patients. Feature selection in NPX data was conducted using three distinct ML methodologies: MUVR (Multivariate Unbiased Variable Reduction), Boruta, and VSURF (Variable Selection Using Random Forests), each contributing unique strengths for optimizing biomarker candidates. MUVR identifies both minimal-optimal variable sets and all-relevant variables, resulting in parsimonious models that minimize variable selection bias and mitigate the risk of model overfitting. The MUVR R package (https://gitlab.com/CarlBrunius/MUVR) was utilized by employing Random Forests (RFs) with 30 repetitions of 30-fold outer cross-validation. All other parameters were maintained at their default settings. The Boruta R package, which also leverages RFs, evaluates feature importance by comparing original variables against randomized shadow features to identify all relevant variables for class prediction. The Boruta (https://cran.r-project.org/web/packages/Boruta/) was used with all parameters set to their default values. Additionally, VSURF was employed as a two-stage approach: it first eliminates irrelevant features and then refines the remaining subset to enhance predictive accuracy through RFs. The VSURF R package (https://cran.r-project.org/web/packages/VSURF) was applied with all parameters maintained at default values.

For the final selection of biomarkers, we focused on the subset of proteins confirmed by at least two out of the three variable selection ML methods. To visualize the overlap among the selected features, a Venn diagram was generated in R using the ggvenn library. This comprehensive approach ensured that the identified biomarkers were reliable and relevant for subsequent analysis. Molecular pathway analysis was conducted using Ingenuity pathway analysis software (IPA) to identify enriched molecular pathways within the protein signature.

### Correlation analysis

Illumina’s Basespace™ Correlation Engine (BSCE; Illumina, San Diego, CA, USA) was used to correlate autoantibodies signature of MS to available MS transcriptome data. BSCE contains over 166,000 Bioset lists of highly curated, statistically significant genes from over 25,100 omic-scale Curated Studies^17^ as of 1st January 2025. Curated Studies in BSCE are processed in standard, platform-specific pipelines to generate gene sets and measures. For each study, available Test group vs. Control group results plus experimental metadata are generated. Second, BSCE has robust data-driven correlation, aggregation, and machine learning applications to exploit the consistent processing and curation of omic-scale studies from international public repositories. The BSCE Disease Atlas application was used to respectively categorize, and rank compounds or diseases based on signature correlations across all BSCE Curated Studies.

## Results

### Study design and cohort specific information

In this study, we evaluated the autoantibody (total IgG) responses in a Qatari-based cohort of Multiple Sclerosis (MS) patients using a high-throughput protein microarray platform comprising 1,600 correctly folded human proteins. The cohort included 39 MS patients and 38 age- and gender-matched healthy controls, all recruited from Hamad Medical Corporation (HMC) in Doha, Qatar (**Supplementary data 1 Figure 1**). While multi-ethnic in composition, the cohort primarily consisted of Qatari nationals. Detailed demographic and clinical characteristics of the participants are summarized in **Table 1**, and **Supplementary Figure 1 and Supplementary data 2** provides an overview of the disease-modifying treatments used by MS patients during the study period. Peripheral blood samples were collected from all participants and processed into serum using a standardized in-house protocol. Serum samples were aliquoted and stored at –80°C until analysis to ensure sample integrity.

**Table 1.**
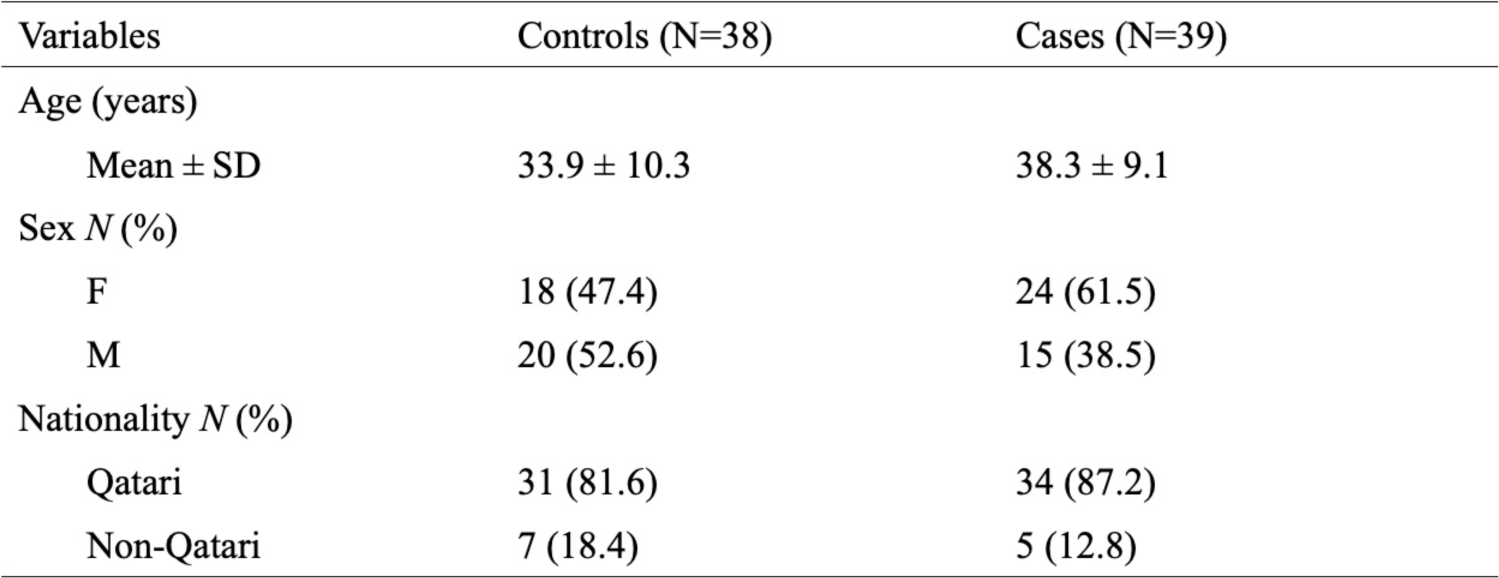

### Elevated Autoantibody Responses to Antiviral and Oligodendrocyte proteins in Multiple Sclerosis

To identify novel functional IgG autoantibodies differentially expressed in Multiple Sclerosis (MS), we employed the high-throughput KoRectly Expressed (KREX) autoantibody assay. This platform enables comprehensive profiling of autoantibody responses against 1,600 human proteins, including kinases, growth factors, interleukins, cytokines, transcription factors, and signaling molecules. Total IgG autoantibody responses were quantified for each antigen using relative fluorescence units (RFU) (**Supplementary Data 3**), which directly correlate with antigen-specific autoantibody titers. Higher RFU values reflect greater concentrations of specific autoantibodies, suggesting elevated levels of their corresponding autoantigens in serum^18, 19^.

Differential expression analysis comparing MS patients and healthy controls revealed significant alterations in autoantibody responses against 129 proteins (**Supplementary Data 4**). Unsupervised hierarchical clustering of these responses clearly distinguished MS patients from controls, as visualized in the heatmap (**Figure 2A**). Volcano plot analysis demonstrated a minimum log2 fold change of ±0.36 and a false discovery rate (FDR) threshold of 0.05, identifying 26 autoantibodies as significantly upregulated and 103 as downregulated in the MS group (**Figure 2B and Supplementary Data 4)**.

**Figure 2.**
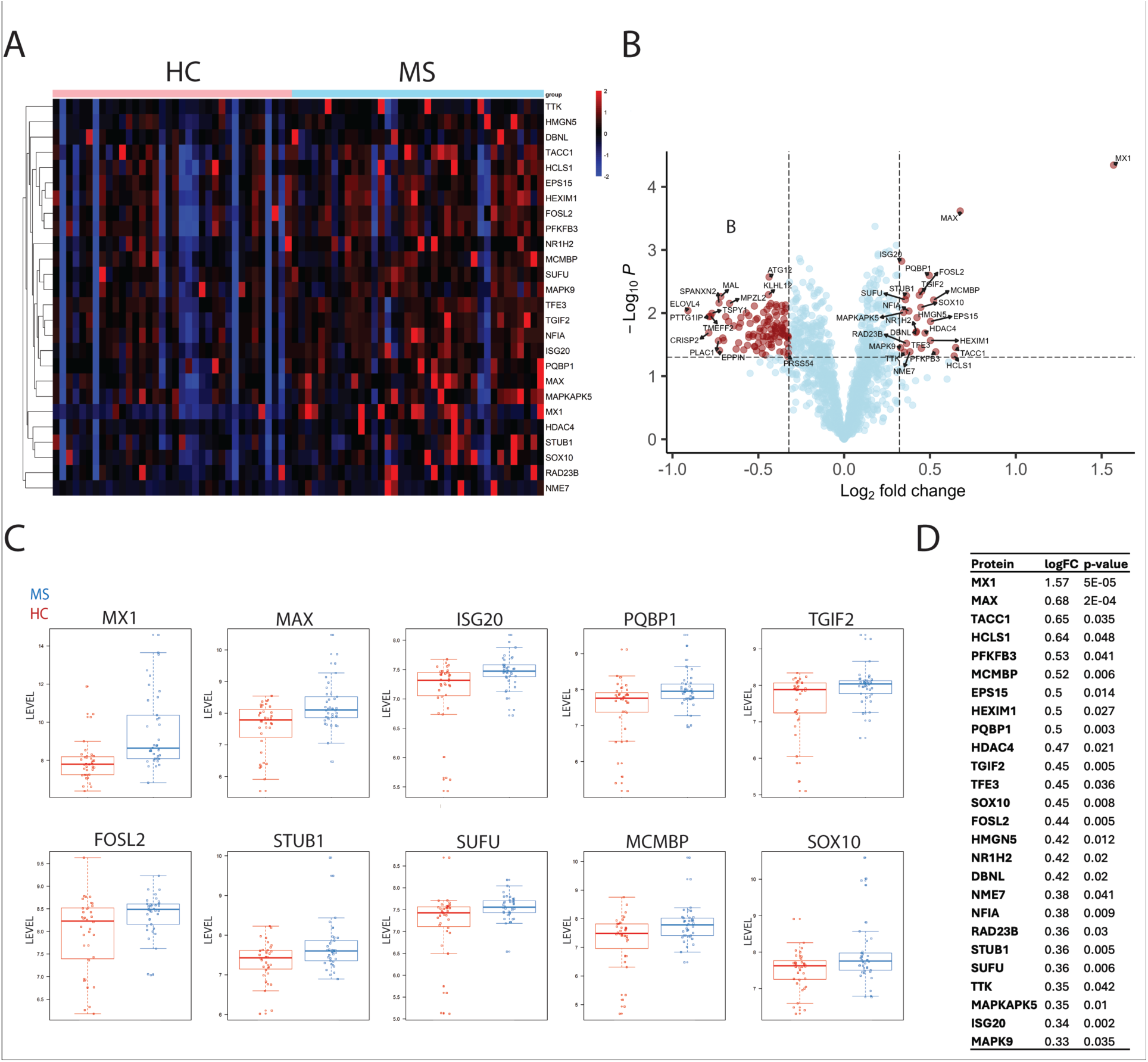

The ten most significantly elevated autoantibodies in MS patients are presented in boxplots and a summary table (**Figure 2C and 2D**). Notably, the most upregulated autoantibody response was directed against Myxovirus resistance protein 1 (MX1), showing a log2 fold change of 1.57 (p < 0.001). MX1 is a well-characterized antiviral protein involved in the type I interferon (IFN-α/β) signaling pathway, previously implicated in MS pathogenesis^20, 21, 22, 23^. Similarly, ISG20— another antiviral immune response protein^24^—also showed elevated autoantibody levels in MS patients.

Intriguingly, we also observed increased autoantibody reactivity against SOX10, a nuclear transcription factor critical for oligodendrocyte differentiation and myelin formation^25, 26, 27^. The presence of autoantibodies against this myelin-associated regulator may indicate ongoing autoimmune responses targeting CNS myelin components.

Conversely, a large subset of autoantibodies (n = 103) were significantly downregulated in MS patients relative to healthy controls (Figure 2B and Supplementary Data), suggesting broader immune dysregulation.

### Machine Learning Reveals a Robust MS Autoantibody Signature

To further elucidate the immunopathology of Multiple Sclerosis (MS), we aimed to define a blood-based autoantibody signature capable of distinguishing MS patients from healthy controls. Using the 129 differentially expressed autoantibodies identified in our initial analysis, we applied three independent machine learning (ML) algorithms—Boruta, MUVR, and VSURF—to identify the most informative biomarkers (**Figure 3, Supplementary Data 5**).

**Figure 3.**
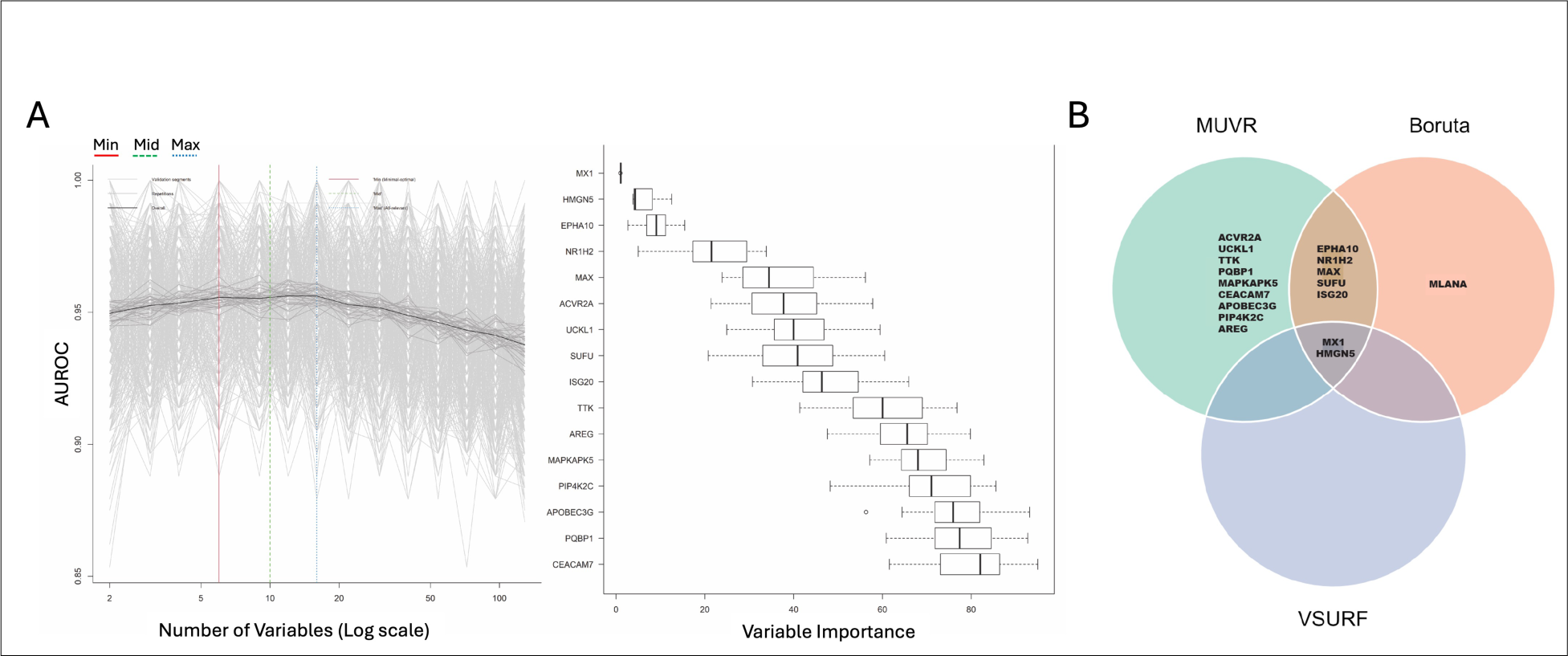

Among these, MUVR yielded the highest discriminatory performance, with a strong area under the receiver operating characteristic curve (AUROC) for its selected biomarkers (**Figure 3A**). A comparative analysis of feature selection across the three ML methods revealed a consistent overlap, notably identifying **MX1** as a top-ranked autoantibody in all models (**Figure 3B**).

This integrative ML approach enabled us to define a robust MS-specific autoantibody signature comprising 17 proteins: **MX1, ISG20, HMGN5, SUFU, MAX, NR1H2, EPHA10, MLANA, ACVR2A, UCKL1, TTK, PQBP1, MAPKAPK5, CEACAM7, APOBEC3G, PIP4K2C**, and **AREG (Figure 3B, Supplementary Data 5)**. These proteins represent a diverse range of functional categories, including antiviral response, transcription regulation, signal transduction, and immune modulation, and may serve as promising candidates for future diagnostic and mechanistic studies in MS.

### Protein Pathway Analysis Reveals Enriched Antiviral Immune and Neurological Signaling pathways in Multiple Sclerosis

To explore the biological relevance of the 17 autoantibody-targeted proteins identified in the MS-specific signature, we conducted Ingenuity Pathway Analysis (IPA). This analysis revealed significant enrichment of pathways involved in immune regulation, antiviral defense, and neurological signaling. Key enriched pathways included interferon alpha/beta signaling, interferon signaling, ISG15 antiviral mechanism, and hypercytokinemia - hyperchemokinemia in the pathogenesis of influenza. In addition, pathways related to the nervous system, such as axonal guidance signaling and EPH-Ephrin signaling, were also prominently represented (**Figure 4, Supplementary Data 5**).

**Figure 4.**
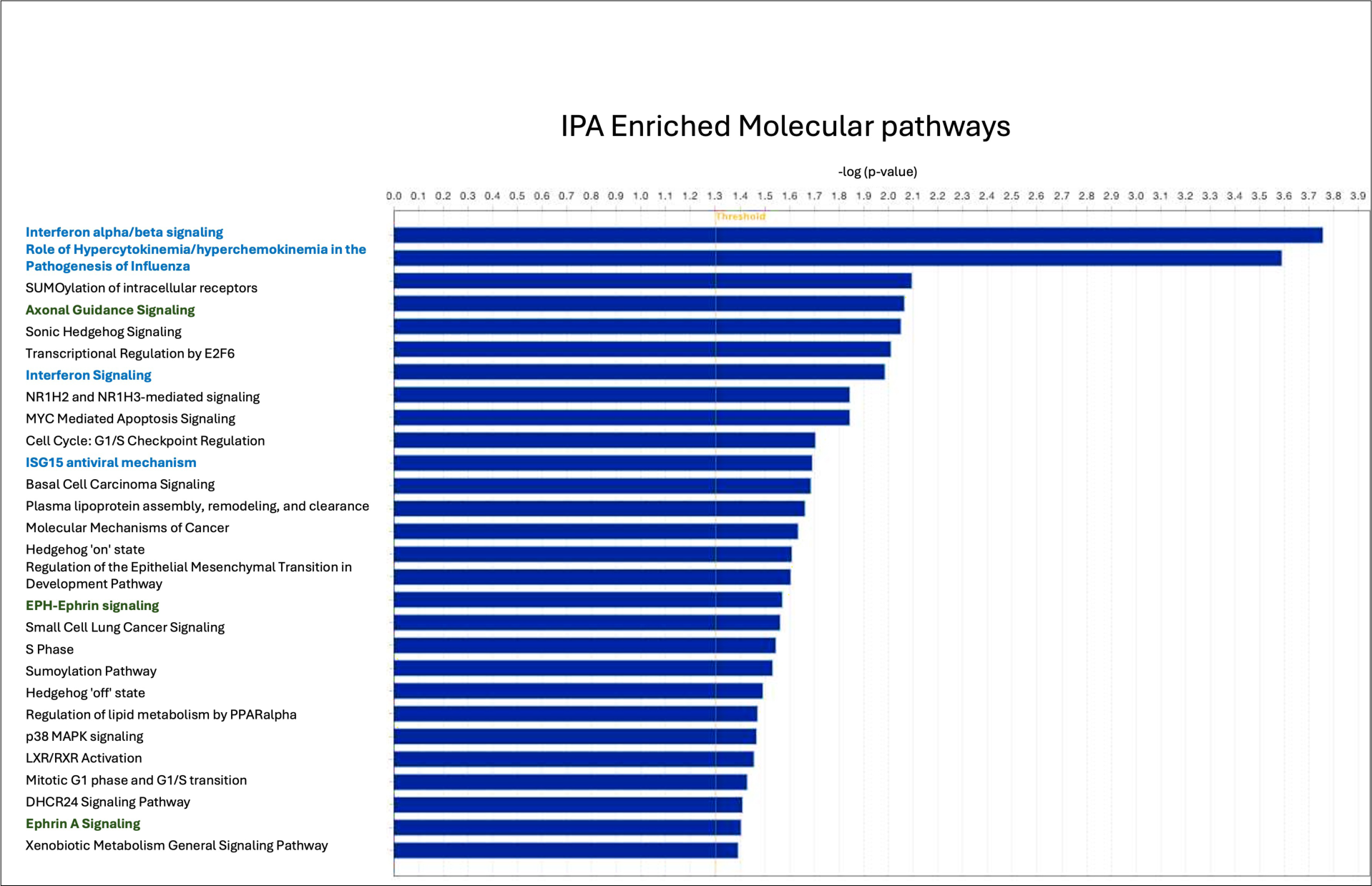

Further IPA-based disease association analysis showed strong links between the MS autoantibody signature and multiple disease categories, including neurological disorders, infectious diseases, ophthalmic conditions, and immunological diseases (**Supplementary Figure 2**). Notably, the protein-protein interaction network derived from the MS autoantibody targets predicted Multiple Sclerosis as an activated disease state (**Supplementary Figure 3**), supporting the biological relevance of these autoantibody markers in MS pathogenesis.

### MS Autoantibody Signature Correlates with Independent Datasets from CNS Injury, Ophthalmic Disease, and Viral Infections

To gain deeper insights into the functional role of the 17-protein MS autoantibody signature, we conducted correlation analyses using transcriptomic data curated in the BSCE Disease Atlas of Multiple Sclerosis. Specifically, we compared our autoantibody signature to RNA expression profiles derived from oligodendrocytes and neurons isolated from chronic inactive MS white matter lesions, which included 5,818 and 8,607 differentially expressed RNAs, respectively (**Figure 5A–B, Supplementary data 6**). In the oligodendrocyte dataset, five overlapping genes were identified: two positively correlated and three negatively correlated with the MS autoantibody profile (**Figure 5A**). Similarly, in the neuron dataset, ten overlapping genes were observed, of which three showed positive correlation and seven showed negative correlation with the signature (**Figure 5B,**).

**Figure 5.**
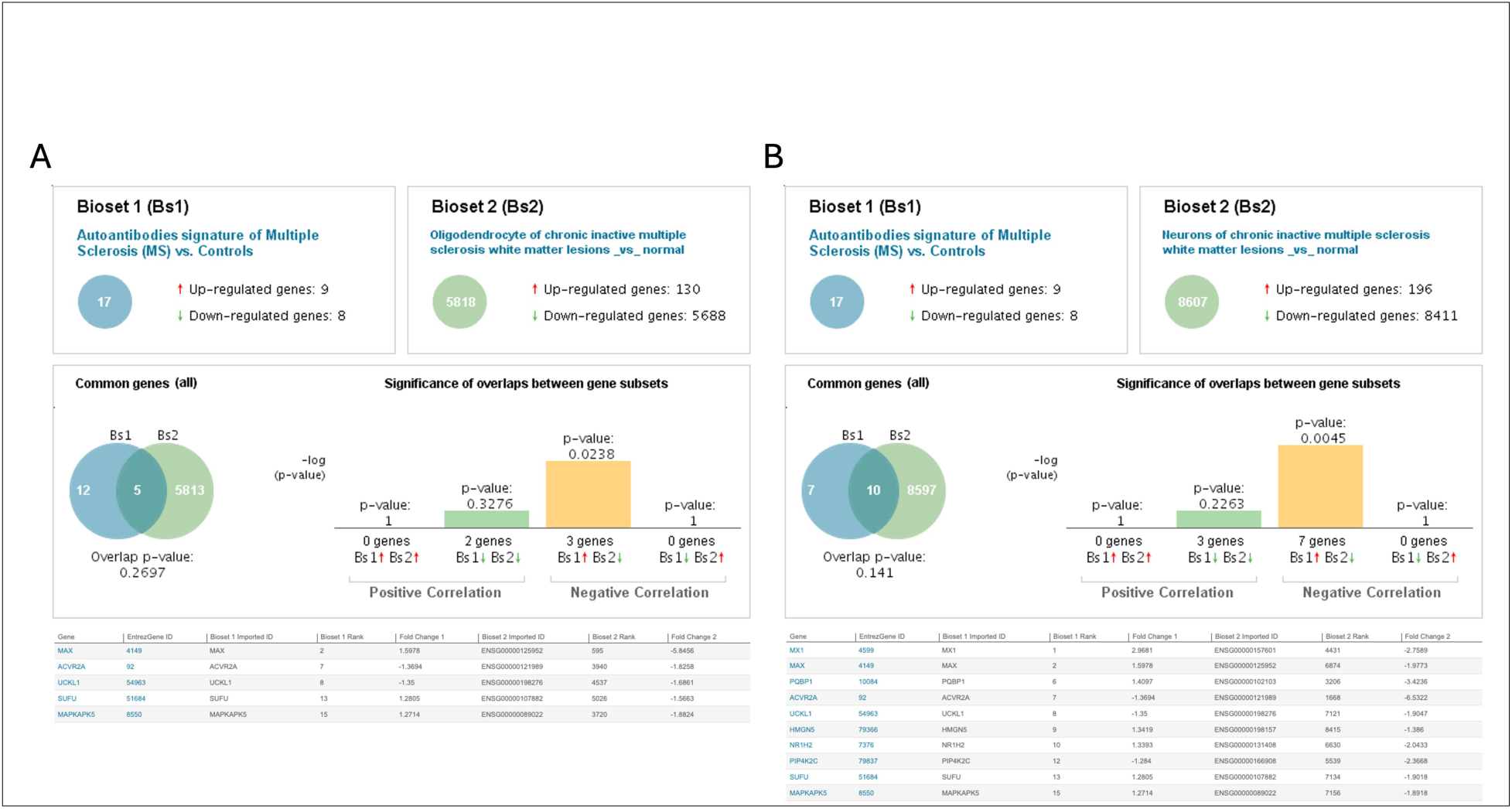
BSCE analysis. Correlation of MS autoantibody signature with transcriptomics data of oligodentdrocytes of chronic inactive multiple sclerosis white matter lesions (A) or neurons of chronic inactive multiple sclerosis white matter lesions (B). Identified proteins with positive and negative correlations are listed in each panel. Red arrow show positive correlation, while green arrow show negative correlation.

To expand these findings, we analyzed transcriptomic data from additional MS-relevant CNS cell types, including microglia, vascular smooth muscle cells, endothelial cells, and pericytes. Across these cell types, most of the overlapping genes correlated negatively with the MS autoantibody signature, with only a few—such as ACVR2A and UCKL1—exhibiting positive correlations (**Table 2**).

**Table 2.**
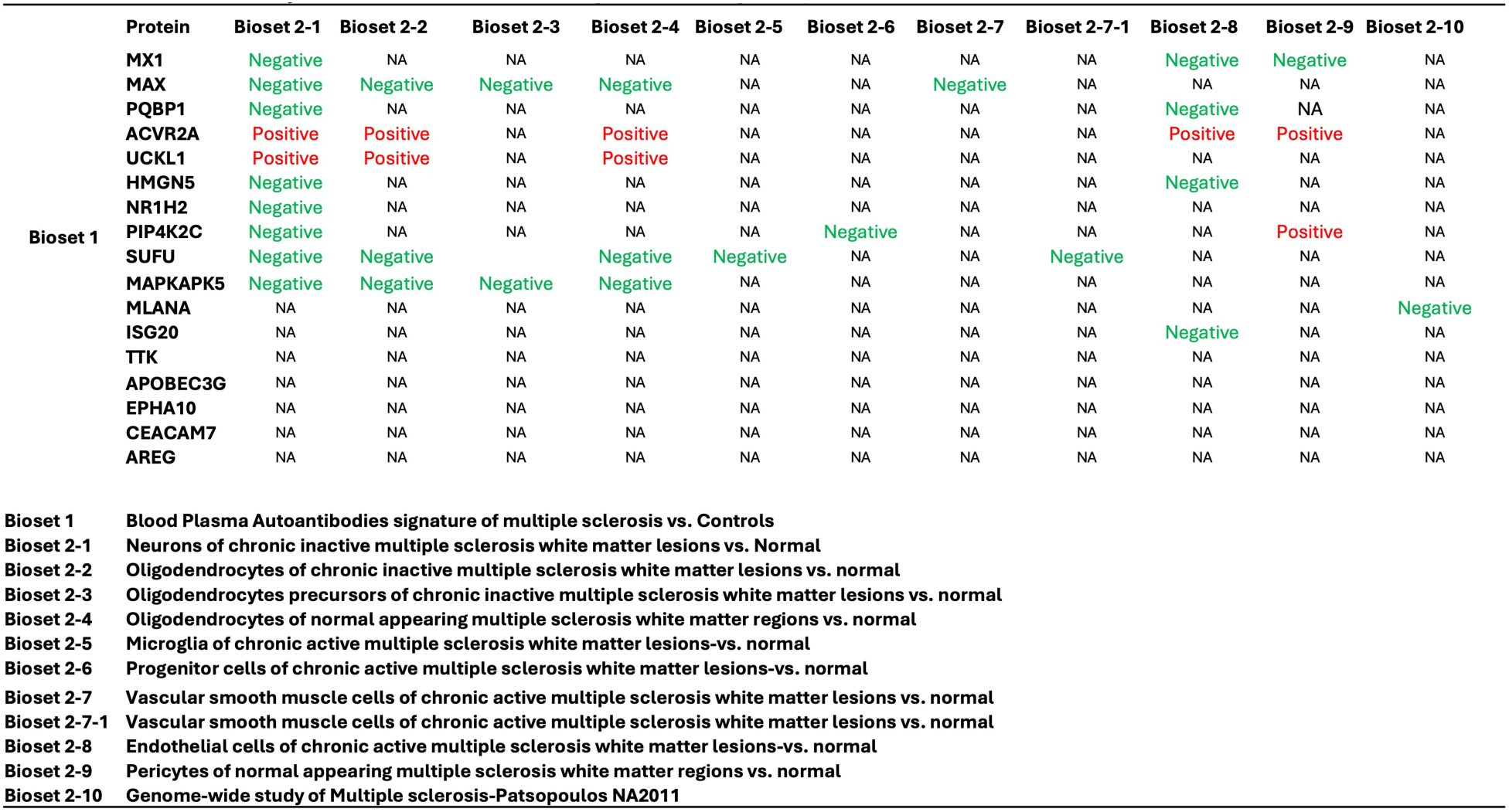
BCCM Correlations analysis between MS autoantibodies signature and MS gene expression data.

We further evaluated the relevance of the MS autoantibody signature in nervous system injuries, using RNA datasets from various animal models. The signature correlated with gene expression changes in spinal cord injury (**Supplementary Figure 4A–B, Supplementary data 6**), dorsal root ganglia after sciatic nerve injury (**Supplementary Figure 4C**), and rat retina following intraorbital nerve crush (**Supplementary Figure 4D**).

Additionally, we assessed correlations with transcriptomic data from ophthalmic disease studies. The MS autoantibody signature showed significant correlations with gene expression profiles from retinal detachment patients (**Supplementary Figure 5A, Supplementary data 6**) and from retina, B cells, and NK cells in age-related macular degeneration (AMD) patients (**Supplementary Figure 5B–D**).

Lastly, we examined potential associations with viral infections using RNA expression data from 24 independent studies (**Supplementary Figure 6, Supplementary data 6**). The MS autoantibody signature displayed both positive and negative correlations across several infections, including Herpes simplex virus (HSV) (**Supplementary Figure 6A–B**) and Cytomegalovirus (CMV) (**Supplementary Figure 6C**), suggesting potential links between past viral exposure and the observed autoimmune responses in MS.

## Discussion

The growing interest in understanding the autoimmune basis of Multiple Sclerosis (MS) continues to drive research aimed at improving diagnosis, monitoring, and treatment of the disease. Despite the availability of immunomodulatory therapies such as interferon-beta (IFN-β), a curative treatment for MS remains elusive. IFN-β, a widely used first-line therapy, offers only partial efficacy, with a significant subset of patients experiencing continued disease activity, including relapses, progression in disability, and formation of new lesions, even under treatment^28–31^.

Recent efforts have focused on the discovery of molecular biomarkers that can inform early diagnosis, predict disease progression, and guide therapeutic decisions. Biomarkers hold promise for minimizing delays in effective treatment and reducing the socioeconomic burden of ineffective drug use^32, 33^. Protein microarrays have emerged as a powerful platform for identifying circulating autoantibodies as potential diagnostic and prognostic biomarkers in MS and other neurodegenerative conditions such as Alzheimer’s and Parkinson’s diseases^34–37^.

In this study, we applied a high-throughput protein microarray (KREX iOme 1600) to identify novel autoantibody responses in a Qatari MS cohort. Machine learning (ML) algorithms were used to isolate a robust MS-specific autoantibody signature composed of 17 key autoantibodies. These proteins are involved in critical immune and neurological pathways, highlighting the multifactorial nature of MS pathophysiology. Notably, we identified MX1 as the most elevated autoantibody in MS patients. While elevated anti-MX1 autoantibodies have been reported in idiopathic pulmonary fibrosis^38^, this is the first report of their involvement in MS. MX1 belongs to the dynamin-like GTPase family and is strongly induced by type I interferons, particularly IFN-α and IFN-β^39,40^. It plays roles in antiviral defense, apoptosis, and cell motility^39,40,41,42^.

Interestingly, MX proteins have been detected in MS brain lesions, particularly in IFN-β–naïve patients, indicating that type I IFN activity may be part of the endogenous immune response in MS^43^. Clinically, MX1 mRNA and protein levels serve as a reliable biomarker of IFN-β bioactivity and treatment response^44, 45, 46, 47^. Reduced MX1 induction after IFN-β treatment is linked to the formation of neutralizing antibodies (NAbs), which inhibit the drug’s efficacy by preventing ISG (interferon-stimulated gene) transcription^46, 48–50^. Thus far, hundreds of ISGs have been identified. They show a strong co-regulation upon stimulation with type I IFNs (IFN-alpha and IFN-beta)^23, 51–54^. Another component of the MS signature, ISG20, is also a type I IFN-stimulated gene and exhibited elevated autoantibody responses in our cohort. Both MX1 and ISG20 are essential in the antiviral IFN pathway and appear to be aberrantly targeted by the immune system in MS patients. Notably, only a small fraction (2.6%) of our cohort received IFN-β treatment (**Sup Figure 1**), suggesting that the elevated immune response to these proteins likely reflects intrinsic disease mechanisms rather than therapy-induced effects. Given their known roles in antiviral immunity, one plausible hypothesis is that viral infections may trigger IFN-mediated upregulation of MX1 and ISG20, inadvertently promoting the development of autoantibodies against these self-proteins. The detection of MX1 and ISG20 autoantibodies in MS patients may impair the therapeutic efficacy of IFN-β as a disease-modifying treatment. Consequently, our findings could support earlier diagnosis and inform treatment strategies prior to the initiation of IFN-β therapy.

MX1 has broad antiviral activity against RNA and DNA viruses^55, 56, 57, 58, 59, 60^, including influenza^61, 62^ and vesicular stomatitis virus^63, 64^. The human cytoplasmic form of MX1 is known to exert broad antiviral effects ^65, 66, 67, 68, 69^. Several studies have provided evidence highlighting Human herpesvirus 6 (HHV-6) and Epstein-Barr virus (EBV) being associated and implicated as etiologic agent in MS^70, 71^. The potential link between viral infections and MS has long been speculated, and our findings offer further support for this hypothesis. We observed positive correlations between the MS autoantibody signature and transcriptomic data from studies involving Herpes simplex virus (HSV) and Cytomegalovirus (CMV) infections. MX1 emerged as a common, positively correlated marker (**Sup. Figure 6**), suggesting that viral activation of IFN pathways could play a role in triggering autoimmunity against IFN-responsive proteins, contributing to MS pathogenesis.

Aberrant immune recognition of self-antigens as foreign agents can lead to the development of autoimmune diseases. This breakdown in immune tolerance may result from persistent infections^72^, continuous antigenic stimulation^73^, impaired mechanisms of self-tolerance^74^, or apoptosis-related processes^75^. The involvement of microbial agents in multiple sclerosis (MS) aligns with its epidemiological patterns and may help explain key aspects of its autoimmune nature. One proposed mechanism is molecular mimicry, in which sequence or structural similarities between microbial components and self-antigens trigger the activation of autoreactive T cells and the production of autoantibodies. For instance, The EBV tegument protein BBRF2, the capsid antigen BFRF3, and EBNA1 exhibit cross-reactivity against CNS autoantigens and MS^76, 77, 78, 79^. Over the past two decades, this hypothesis has gained considerable support^71, 80, 81, 82, 83^. Driven by both epidemiological evidence and the appeal of the mimicry model, numerous studies have explored the presence of viruses in the cerebrospinal fluid (CSF) and other biological samples from MS patients^70^. Multiple viruses have been implicated in the disease. Therefore, reviewing the medical history of MS patients for prior viral infections may provide valuable insight into the onset and progression of MS. In addition to MX1, other autoantibodies identified in MS that show either positive or negative correlations with viral infection data warrant detailed investigation.

Given the autoimmune nature of multiple sclerosis (MS), substantial research has focused on identifying immune-related biomarkers that could facilitate early diagnosis. Autoantibodies targeting myelin surface proteins—such as myelin oligodendrocyte glycoprotein (MOG), myelin basic protein (MBP), proteolipid protein (PLP), and myelin-associated glycoprotein (MAG)— have shown associative or correlative links to MS. However, these antibodies currently lack the specificity and reliability required for diagnostic application^84, 85, 86, 87, 88, 89, 90^. Other emerging autoantibody targets include glycans like GAGA4 and proteins such as KIR4.1, which are drawing increasing attention^91, 34, 92, 93, 94^.

Recent studies have revealed that MS lesions exhibit a heterogeneous gene expression profile related to oligodendrogliogenesis. Notably, the transcription factor SOX10, which is essential for oligodendrocyte maturation and the myelination program^25, 26, 27, 95^, is significantly downregulated in inactive demyelinated MS lesions compared to normal-appearing white matter^96^. Building on this, García-León and colleagues demonstrated that SOX10 overexpression in human pluripotent stem cells (hPSCs) derived from MS patients is sufficient to induce differentiation into O4-positive (O4⁺) and MBP-positive oligodendrocytes^97, 98^. Strikingly, in our MS cohort, we observed elevated levels of autoantibodies targeting SOX10 (**Figure 2C**). This immune response could potentially interfere with oligodendrocyte-mediated remyelination in MS lesions^95, 99^. Further research is warranted to elucidate the role of SOX10 autoantibodies in MS pathogenesis, including their association with clinical phenotypes, disease progression, and severity.

Since MS is a cell type-specific disease that primarily targets oligodendrocytes in the CNS, it is crucial to investigate whether the autoantibodies we detected in serum are also present in the cerebrospinal fluid (CSF), and whether viral infections and activation of the interferon-beta (IFN-β) pathway—particularly through MX1 and ISG20—are linked to oligodendrocyte involvement.

Overall, these findings emphasize the complexity of MS as a disease involving both immune dysregulation and neurodegenerative processes. While previous efforts have focused on myelin-related autoantigens, newer targets like SOX10 and immune pathway proteins like MX1 and ISG20 provide deeper insights into disease mechanisms and may represent novel therapeutic or diagnostic avenues.

## Conclusions

In summary, our study uncovers a distinct autoantibody response profile in MS patients, targeting proteins involved in both immune regulation and neurological function. These findings emphasize the significant role of the humoral immune response and suggest a potential link between prior viral infections and MS pathogenesis. Notably, elevated autoantibody levels against MX1 and ISG20 point to novel molecular associations that may contribute to disease development and progression. Future research directions should aim to 1) Validate the Qatar-derived MS autoantibody signature in independent, multiethnic cohorts using orthogonal assays and functional models; 2) Investigate the mechanistic impact of these autoantibodies on pathways involved in MS onset, activity, severity, and progression; and 3) Correlate the identified autoantibody biomarkers with clinical MS subtypes and disease-modifying therapies, and patient outcomes. These efforts will be critical for advancing our understanding of MS immunopathology and may support the development of novel diagnostic and therapeutic strategies.

## Author Contributions

H.B.A designed, conceived, and led the study. I.N.P., and R.A.M., provided the cohort samples and clinical data. H.B.A., and I.B. ran the assays. D.A. and T.M.T performed Sengenics data preprocessing and quality control analyses, B.A., L.K., supported Sengenics study design and management. A.D.V.B. performed bioinformatics data analyses. H.B.A performed IPA and BSCE analyses. H.B.A., I.N.P., and R.A.M., interpreted the data. H.B.A wrote the manuscript. All authors reviewed the manuscript and have read and agreed to the published version of the manuscript.

## Institutional Review Board Statement

The study on Qatar Cohort was conducted according to the Ministry of Public Health (MOPH) guidelines and approved by the Institutional Review Board Research Ethics Committee of Weill Cornell Medicine Qatar (reference IRB # 23-00031).

## Acknowledgments

We extend our sincere gratitude to all the patients from Hamad Medical Corporation for their participation. We also acknowledge the Qatar Biobank for providing multiple sclerosis and control samples, as well as for their support in sample selection and transfer to the HBKU Core Lab Facility. This study was supported by the Proteomics Core Laboratory at Hamad Bin Khalifa University.

## Glossary

ACVR2A: Activin A receptor type 2A
APOBEC3G: Apolipoprotein B mRNA editing enzyme catalytic subunit 3G
AREG: Amphiregulin
CEACAM7: CEA cell adhesion molecule 7
CMV: Cytomegalovirus
CNS: Central Nervous System
CSF: CerebroSpinal Fluid
DEPs: Differentially Expressed Proteins
EBV: Epstein–Barr virus
EPHA10: Ephrin receptor A10
HMGN5: High mobility group nucleosome binding domain 5
HSV: Herpes simplex virus
INF: Interferon
IPA: Ingenuity Pathway Analysis
ISG20: Interferon stimulated exonuclease gene 20
ISGs: Interferon-stimulated genes
KREX: KoRectly Expressed
MAPKAPK5: MAPK activated protein kinase 5
MAX: MYC associated factor X
MLANA: Melan-A
MS: Multiple sclerosis
MX1: Myxovirus resistance protein 1
NAbs: Neutralizing antibodies
NK: Natural killer
NPX: Normalized Protein Expression
NR1H2: Nuclear receptor subfamily 1 group H member 2
PIP4K2C: Phosphatidylinositol-5-phosphate 4-kinase type 2 gamma
PQBP1: Polyglutamine binding protein 1
RFU: Relative Fluorescence Unit
SOX-10: SRY-box transcription factor 10
SUFU: Suppressor of Fused, negative regulator of the Hedgehog (HH) signalling pathway
TTK: Threonine tyrosine kinase
UCKL1: Uridine-cytidine kinase 1 like 1

---

**Sup Figure 1.**
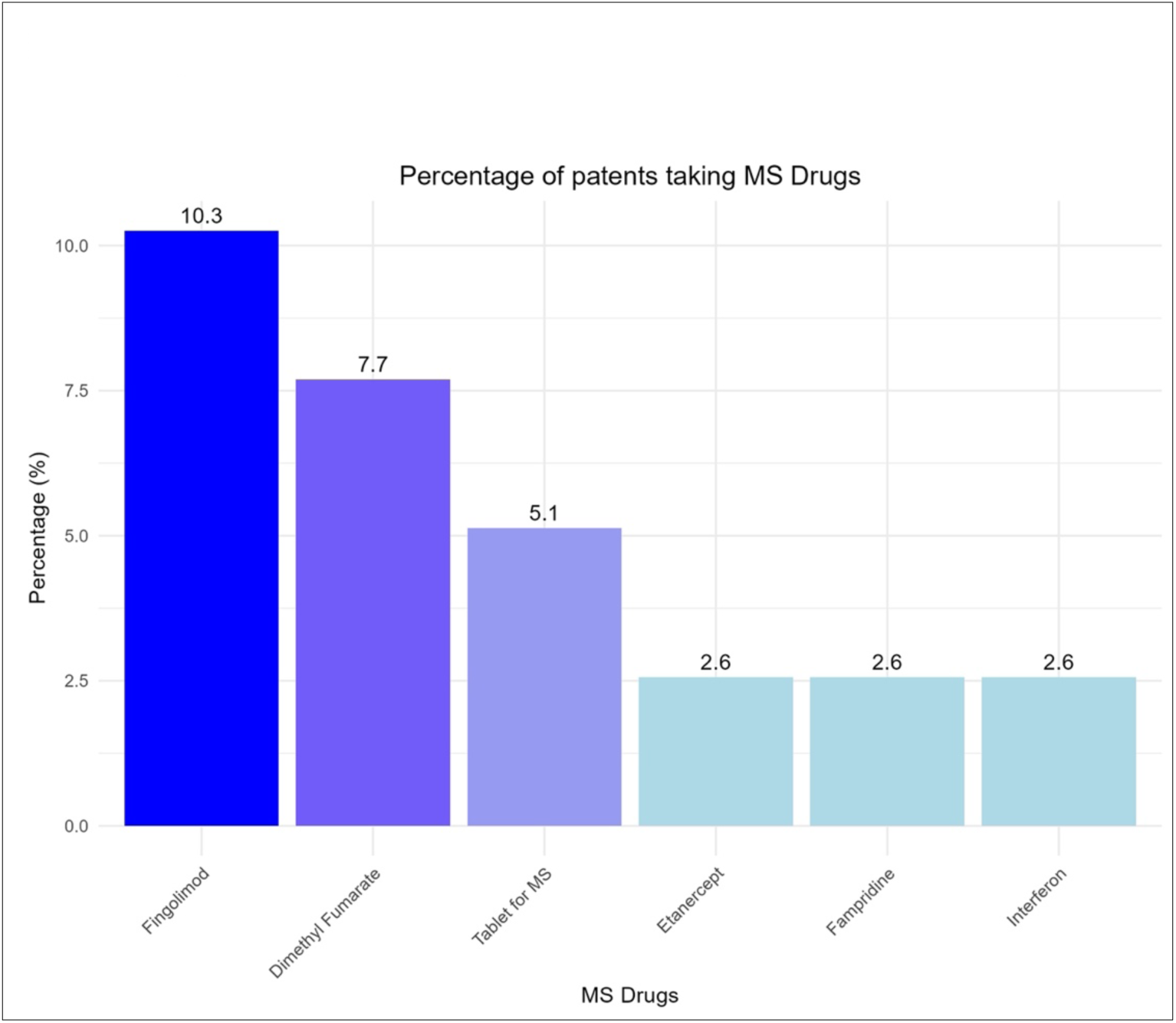

**Sup Figure 2.**
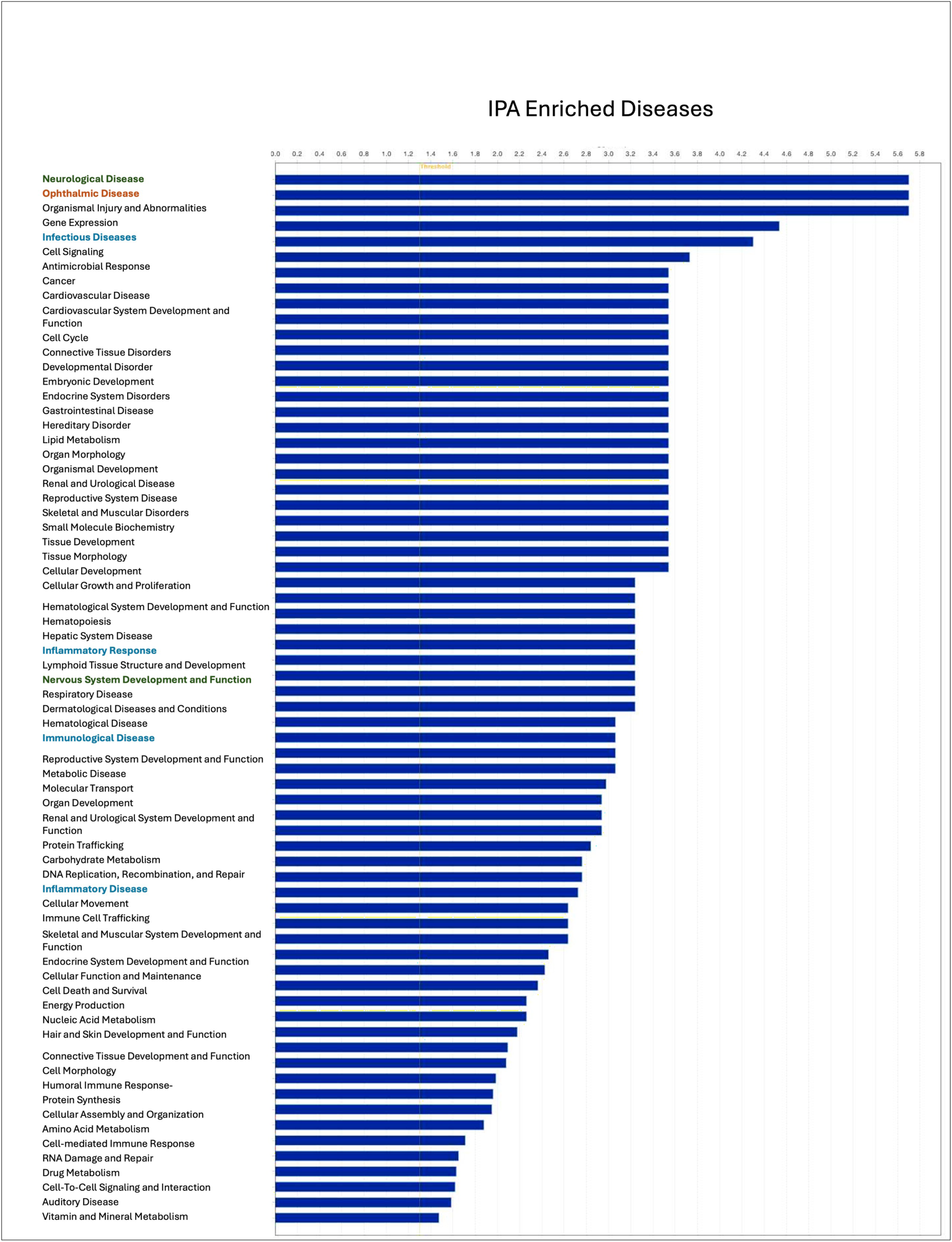

**Sup Figure 3.**
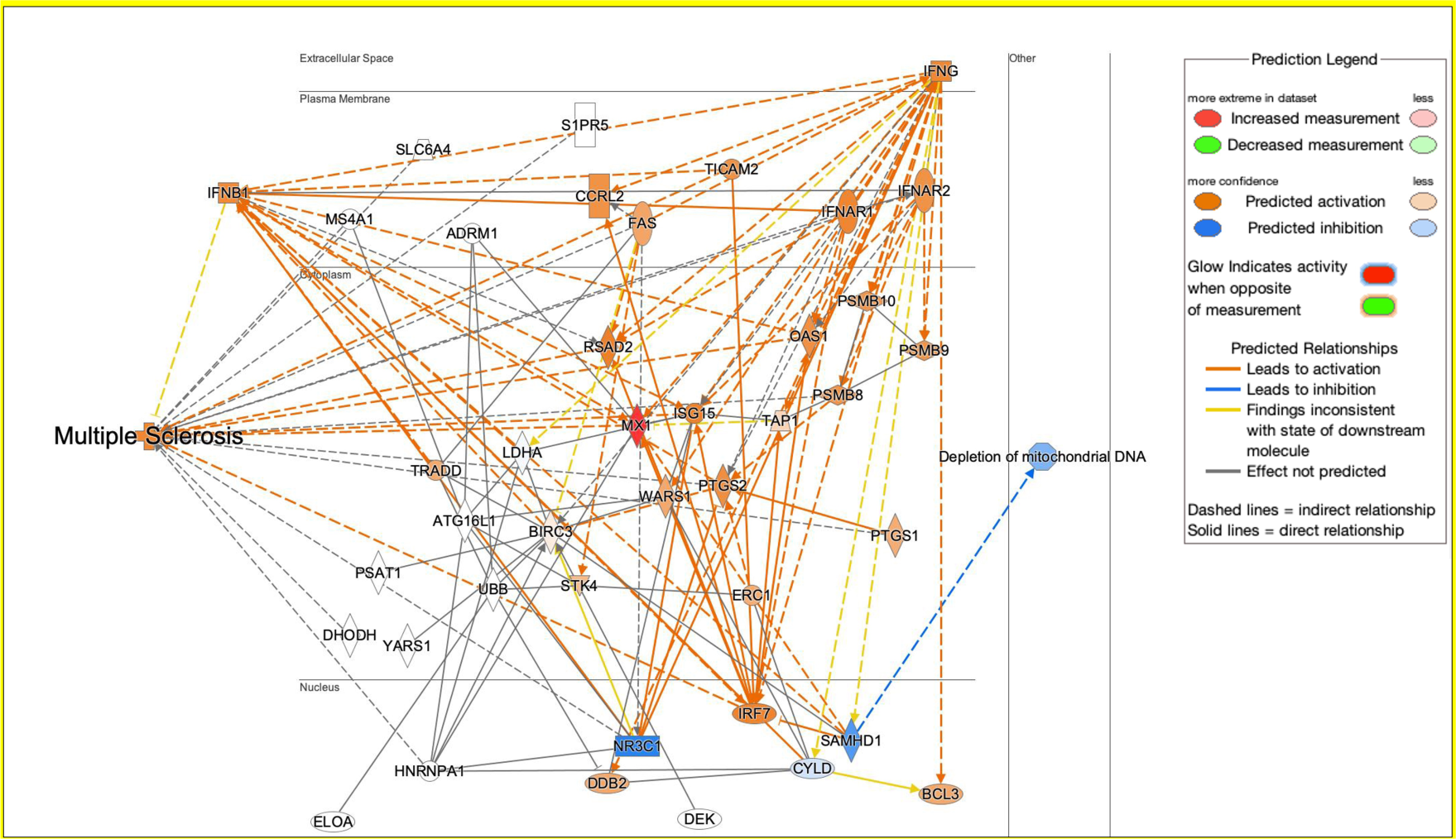

**Sup Figure 4.**
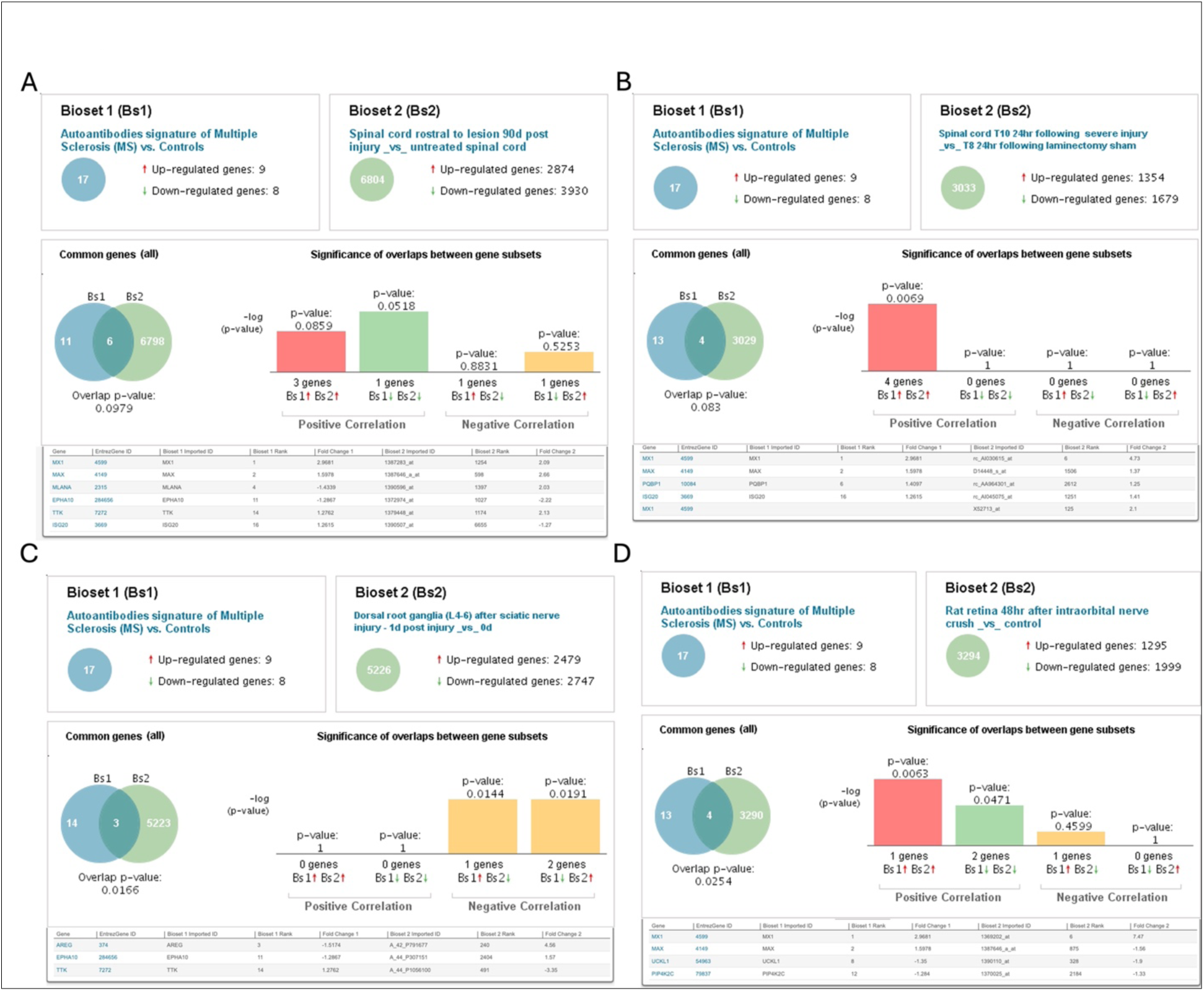
BSCE analysis. Correlation of MS autoantibody signature with transcriptomics data of (A) or (B). Identified proteins with positive and negative correlations are listed in each panel. Red arrow show positive correlation, while green arrow show negative correlation.

**Sup Figure 5.**
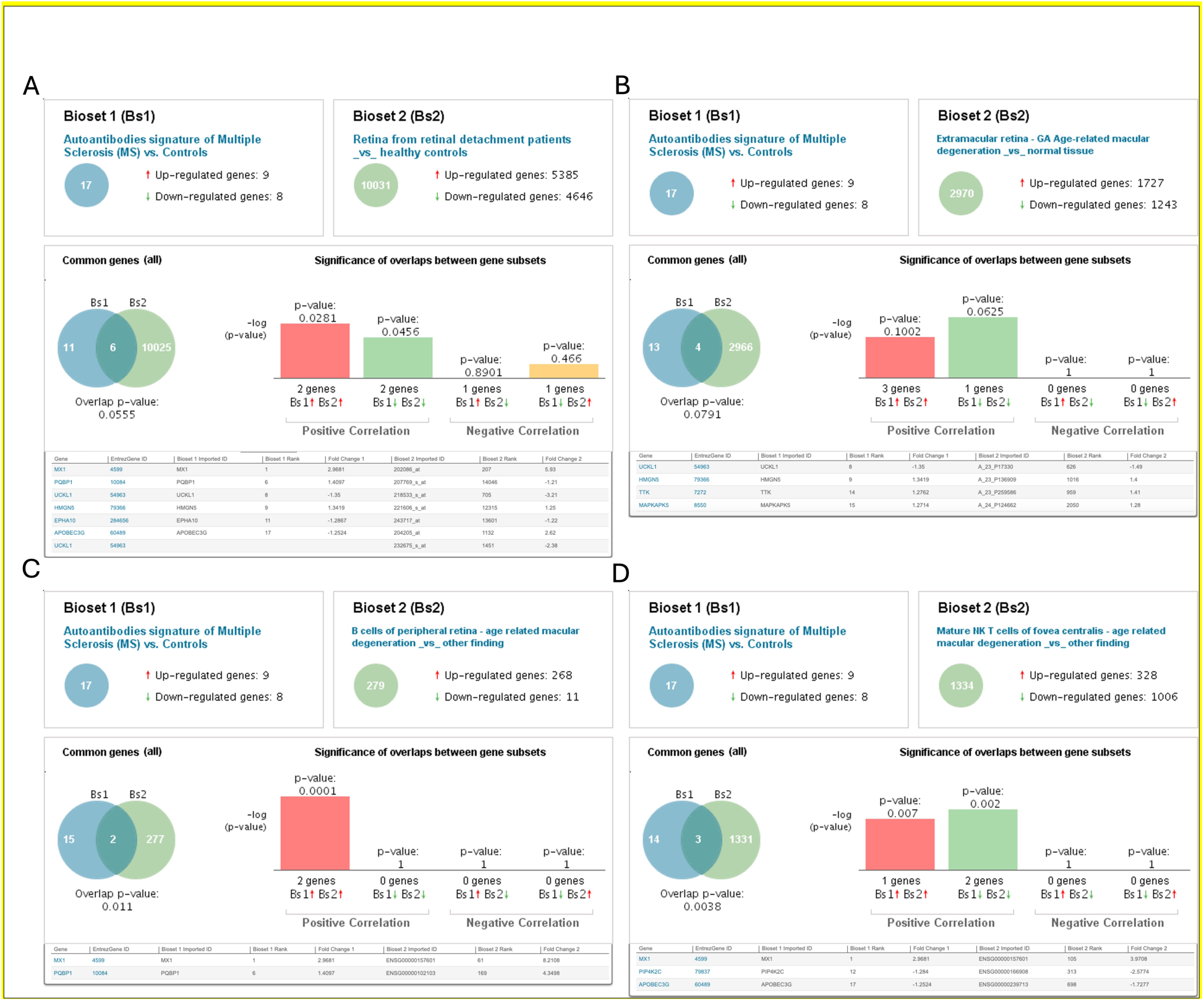
BSCE analysis.Correlation of MS autoantibody signature with transcriptomics data of (A) or (B). Identified proteins with positive and negative correlations are listed in each panel. Red arrow show positive correlation, while green arrow show negative correlation.

**Sup Figure 6.**
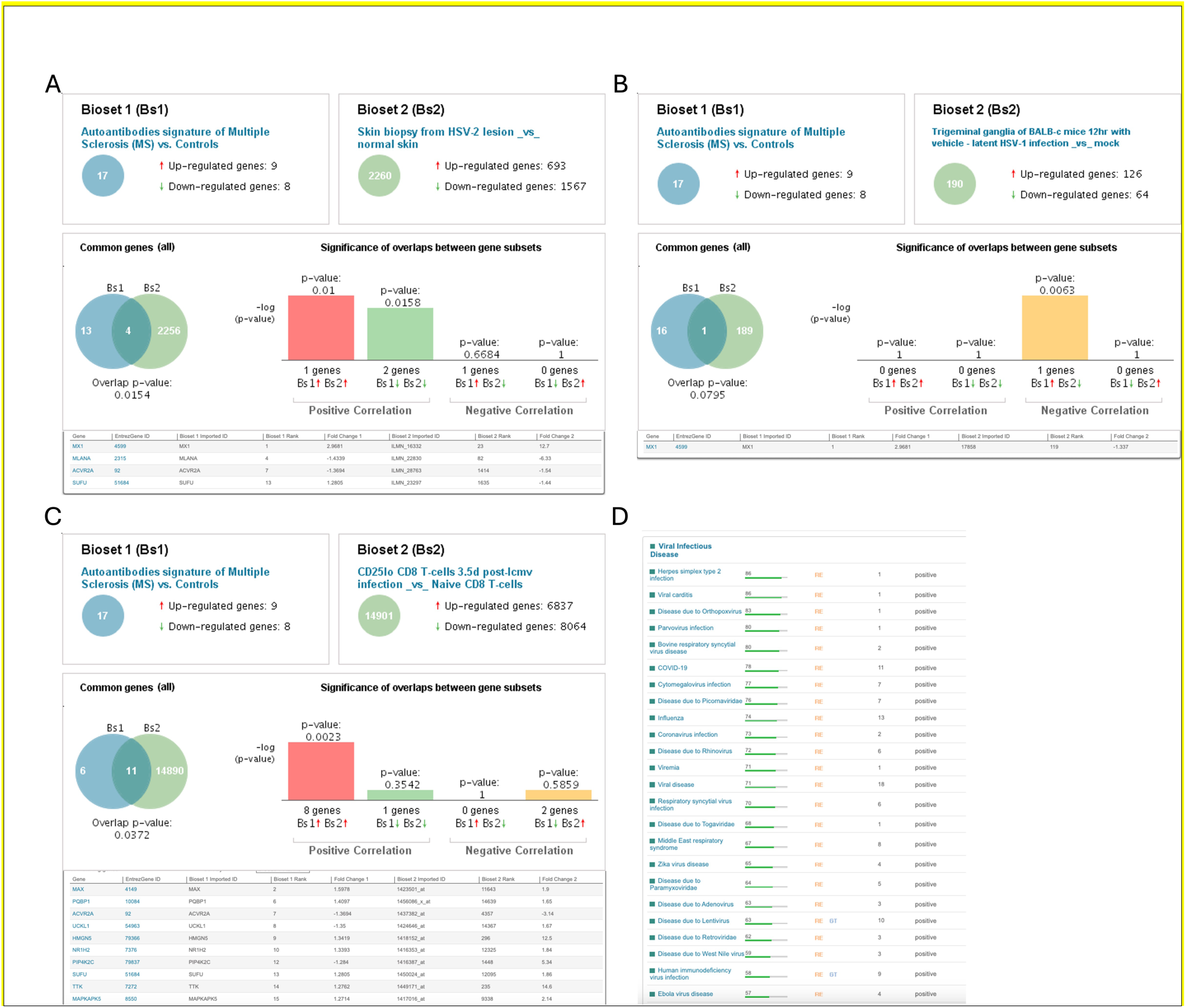
BSCE analysis. Correlation of MS autoantibody signature with transcriptomics data of (A) or (B). Identified proteins with positive and negative correlations are listed in each panel. Red arrow show positive correlation, while green arrow show negative correlation.

