## Supplementary material for "A New Serological Autoantibody Signature Associated with Multiple Sclerosis": sheet001.htm

|  |  |  |  |  |  |  |
| --- | --- | --- | --- | --- | --- | --- |
| Project\_Dummy\_ID | Case/ Control | Gender | Nationality | Age | HEIGHTWEIGHT\_OUT\_WEIGHT | BMI |
| WC000268000001 | Control | Female | Qatari | 22 | 84.9 | 32.96 |
| WC000268000002 | Control | Female | Non-Qatari | 48 | 85.8 | 34.33 |
| WC000268000003 | Control | Female | Qatari | 28 | 67.25 | 27.6 |
| WC000268000004 | Control | Female | Non-Qatari | 40 | 77.6 | 31.56 |
| WC000268000005 | Control | Female | Qatari | 48 | 90.95 | 33.65 |
| WC000268000006 | Control | Female | Qatari | 22 | 73.75 | 26.22 |
| WC000268000007 | Control | Female | Qatari | 21 | 50 | 21.17 |
| WC000268000008 | Control | Female | Non-Qatari | 39 | 88.15 | 33.3 |
| WC000268000009 | Control | Female | Qatari | 25 | 65.35 | 25.4 |
| WC000268000010 | Control | Female | Qatari | 36 | 63.1 | 25.18 |
| WC000268000011 | Control | Female | Qatari | 42 | 69.3 | 27.62 |
| WC000268000012 | Control | Female | Qatari | 19 | 49.6 | 19.74 |
| WC000268000013 | Control | Female | Qatari | 45 | 81.2 | 32.94 |
| WC000268000014 | Control | Female | Qatari | 28 | 65.2 | 22.99 |
| WC000268000015 | Control | Male | Qatari | 51 | 95.3 | 35.56 |
| WC000268000017 | Control | Female | Qatari | 25 | 49 | 22.07 |
| WC000268000018 | Control | Female | Qatari | 18 | 51.8 | 22.63 |
| WC000268000019 | Control | Female | Qatari | 27 | 62.45 | 23.83 |
| WC000268000020 | Control | Female | Qatari | 18 | 105.05 | 34.94 |
| WC000268000022 | Control | Male | Non-Qatari | 38 | 102.7 | 32.52 |
| WC000268000047 | Control | Male | Qatari | 61 | 95.95 | 33.63 |
| WC000268000048 | Control | Male | Non-Qatari | 32 | 71.95 | 23.02 |
| WC000268000050 | Control | Male | Qatari | 45 | 75.85 | 28.62 |
| WC000268000051 | Control | Male | Qatari | 28 | 77.5 | 27.62 |
| WC000268000052 | Control | Male | Non-Qatari | 39 | 74.5 | 24.49 |
| WC000268000053 | Control | Male | Qatari | 29 | 95.85 | 33.28 |
| WC000268000055 | Control | Male | Non-Qatari | 40 | 117.4 | 38.33 |
| WC000268000056 | Control | Male | Qatari | 43 | 82.05 | 28.8 |
| WC000268000058 | Control | Male | Qatari | 31 | 82.85 | 24.96 |
| WC000268000059 | Control | Male | Qatari | 32 | 82.3 | 30.27 |
| WC000268000064 | Control | Male | Qatari | 33 | 80 | 28.82 |
| WC000268000065 | Control | Male | Qatari | 35 | 120.2 | 38.54 |
| WC000268000066 | Control | Male | Qatari | 18 | 84.15 | 28.68 |
| WC000268000067 | Control | Male | Qatari | 40 | 100.5 | 32.04 |
| WC000268000068 | Control | Male | Qatari | 46 | 81.45 | 27.89 |
| WC000268000072 | Control | Male | Qatari | 31 | 82.8 | 28.09 |
| WC000268000073 | Control | Male | Qatari | 31 | 91.4 | 32.19 |
| WC000268000076 | Control | Male | Qatari | 35 | 62.55 | 23.43 |
| WC000268000016 | Case | Female | Qatari | 46 | 55.6 | 24.13 |
| WC000268000021 | Case | Female | Non-Qatari | 31 | 54.6 | 24.23 |
| WC000268000024 | Case | Female | Qatari | 34 | 81.1 | 30.49 |
| WC000268000025 | Case | Female | Qatari | 33 | 84.8 | 29.73 |
| WC000268000026 | Case | Female | Qatari | 42 | 79.25 | 29.94 |
| WC000268000027 | Case | Female | Qatari | 39 | 69.4 | 27.59 |
| WC000268000028 | Case | Female | Qatari | 44 | 74.7 | 25.79 |
| WC000268000029 | Case | Female | Qatari | 35 | 64.1 | 29.38 |
| WC000268000030 | Case | Female | Qatari | 48 | 68 | 29.09 |
| WC000268000031 | Case | Female | Qatari | 24 | 78.85 | 29.03 |
| WC000268000032 | Case | Female | Qatari | 37 | 78.75 | 32.11 |
| WC000268000033 | Case | Female | Qatari | 37 | 54.9 | 24.8 |
| WC000268000034 | Case | Female | Qatari | 39 | 72.25 | 28.72 |
| WC000268000035 | Case | Female | Qatari | 24 | 56.55 | 22.23 |
| WC000268000036 | Case | Female | Qatari | 41 | 82.05 | 34.51 |
| WC000268000037 | Case | Female | Qatari | 75 | 71.3 | 31.02 |
| WC000268000038 | Case | Female | Qatari | 42 | 109.8 | 40.33 |
| WC000268000039 | Case | Female | Qatari | 33 | 70.65 | 29.14 |
| WC000268000040 | Case | Female | Qatari | 33 | 69.7 | 23.61 |
| WC000268000041 | Case | Female | Qatari | 33 | 63.5 | 23.64 |
| WC000268000042 | Case | Female | Qatari | 38 | 83.15 | 31.88 |
| WC000268000043 | Case | Female | Qatari | 40 | 87 | 31.38 |
| WC000268000044 | Case | Female | Qatari | 47 | 78.8 | 31.61 |
| WC000268000045 | Case | Male | Qatari | 38 | 93.3 | 28.42 |
| WC000268000046 | Case | Female | Qatari | 44 | 57.2 | 22.37 |
| WC000268000049 | Case | Male | Qatari | 35 | 77.6 | 29.1 |
| WC000268000054 | Case | Male | Non-Qatari | 27 | 88.55 | 28.04 |
| WC000268000057 | Case | Male | Qatari | 28 | 105.95 | 32.3 |
| WC000268000060 | Case | Male | Qatari | 34 | 71.2 | 22.07 |
| WC000268000061 | Case | Male | Qatari | 43 | 79.75 | 29.26 |
| WC000268000062 | Case | Male | Qatari | 44 | 72.65 | 25.93 |
| WC000268000063 | Case | Male | Non-Qatari | 48 | 99 | 31.96 |
| WC000268000069 | Case | Male | Qatari | 37 | 100.55 | 34.96 |
| WC000268000070 | Case | Male | Qatari | 34 | 75.25 | 25.06 |
| WC000268000071 | Case | Male | Qatari | 45 | 107.75 | 31.72 |
| WC000268000074 | Case | Male | Qatari | 42 | 163.55 | 52.32 |
| WC000268000075 | Case | Male | Non-Qatari | 45 | 109.1 | 35.58 |
| WC000268000077 | Case | Male | Non-Qatari | 24 | 54.55 | 17.63 |
| WC000268000078 | Case | Male | Qatari | 29 | 85.95 | 27.87 |
