## Supplementary material for "A New Serological Autoantibody Signature Associated with Multiple Sclerosis": sheet002.htm

|  |  |  |  |  |
| --- | --- | --- | --- | --- |
| Variables | Controls | Cases |  |  |
| N | 38 | #REF! |  | |
| Age (Years) | 33.9 | 38.3 |  | |
| Mean (±SD) | 10.3 | 9.1 |  | |
| Sex(N) |  | | % | % |
| F | 18 | 24 | 47.4 | #REF! |
| M | 20 | 15 | 52.6 | #REF! |
| Nationality (N) |  | | % | % |
| Qatari | 31 | 34 | 81.6 | #REF! |
| Non-Qatari | 7 | 5 | 18.4 | #REF! |
| Weight (±SD) | 79.9 | 80.3 |  | |
|  | 17.3 | 20.6 |  | |
| BMI (±SD) | 28.9 | 29.2 |  | |
|  | 4.9 | 5.8 |  | |
